## Supplementary figures for "IdentifiHR: predicting homologous recombination deficiency in high-grade serous ovarian carcinoma using gene expression"

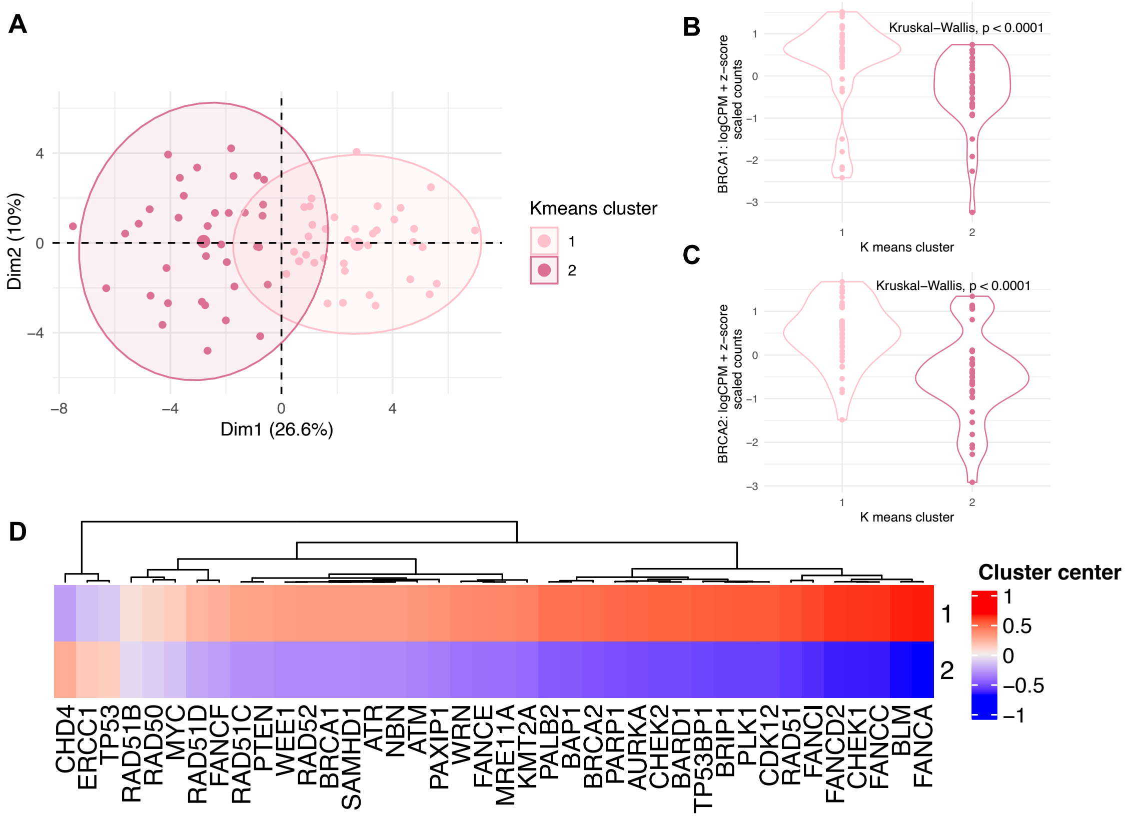


**Supplementary figure 1.** *K means clustering of the BRCAness gene expression signature in the TCGA testing cohort (n = 73)*. (A) Counts of all genes were $log_2$ CPM transformed, then subset to only genes within the BRCAness signature and z score scaled before being used as input for k means clustering, with 2 centroids. The expression of HR associated genes of the signature, specifically (B) *BRCA1* and (C) *BRCA2* were examined by k means clusters in violin plots, where each point represents a sample, with Kruskal-Wallis testing for expression differences, where significance is defined at $p < 0.05$. (D) The cluster centers of each gene within the signature were also examined to determine the HR status best represented by each cluster.

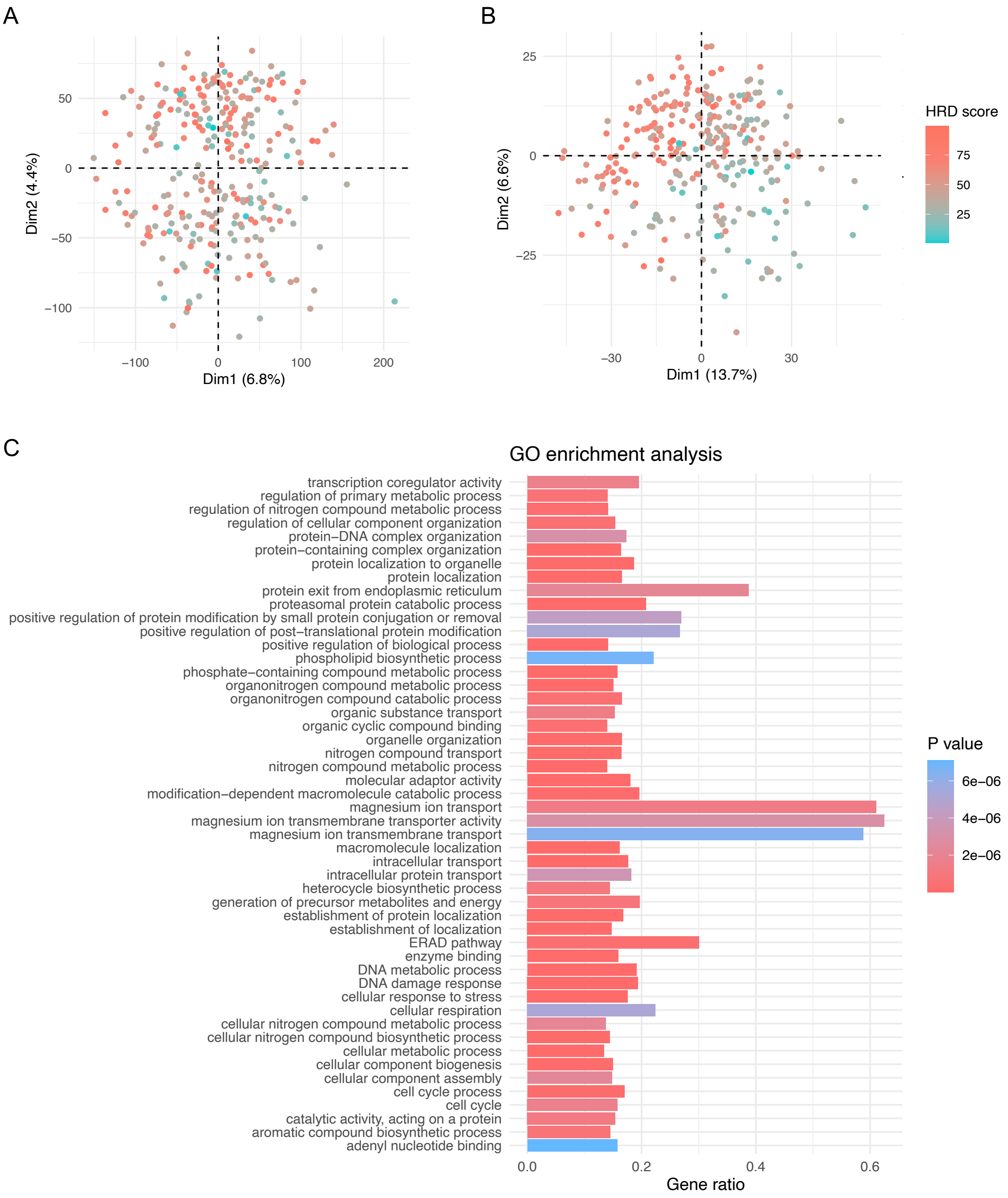


**Supplementary figure 2.** *Significantly DE genes were used for IdentifiHR feature selection to predict HR status*. PCA plots of the first and second principal components after counts of (A) all genes (n = 52125) and (B) DE genes (n = 2604) were normalised by $\log_2$ CPM and scaled using a z-score. Each point represents a unique sample of the TCGA training cohort (n = 288), coloured by HRD score. (C) The top 50 gene ontologies significantly enriched for differentially expressed genes between HRD and HRP HGSCs. 'Gene ratio' shows the number of DE genes divided by the total number of genes in the ontology set.

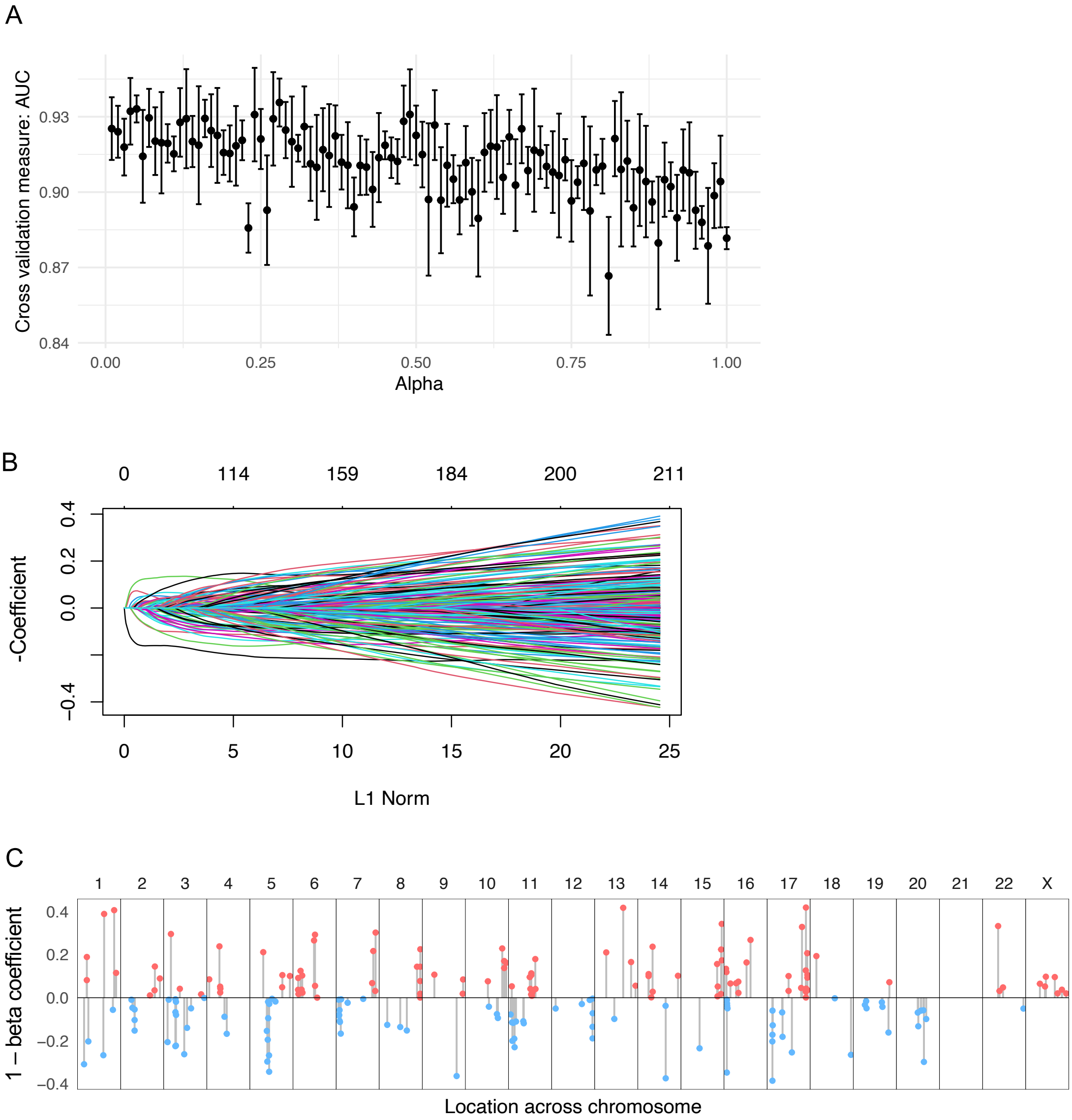


**Supplementary figure 3.** *209 features were included in the IdentifiHR model, following hyperparameter tuning of alpha and lambda*. (A) Selection of the optimal alpha hyperparameter value, with the associated AUC taken from 5-fold cross validation shown by points, with error bars of 1 standard deviation from each AUC. An alpha of 0.53 was selected, being the largest alpha value (in the sequence from 0.1 to 1.0, in intervals of 0.1) within 1 standard deviation of the highest AUC for any alpha value. The resulting model required 209 genes as input features; (B) beta coefficients of these features were symmetrically weighted, given by coloured lines, each of which represents a gene in the model. Number of genes at each level of L1 norm given by integers at the top of the plot. (C) Chromosomal locations of the 209 model genes given as stemmed points, by the associated 1- beta coefficient that weights the gene in the model. The inverse beta coefficient was visualised for easier comparison with differential expression analysis results. Chromosomes lengths are scaled and given by black borders.

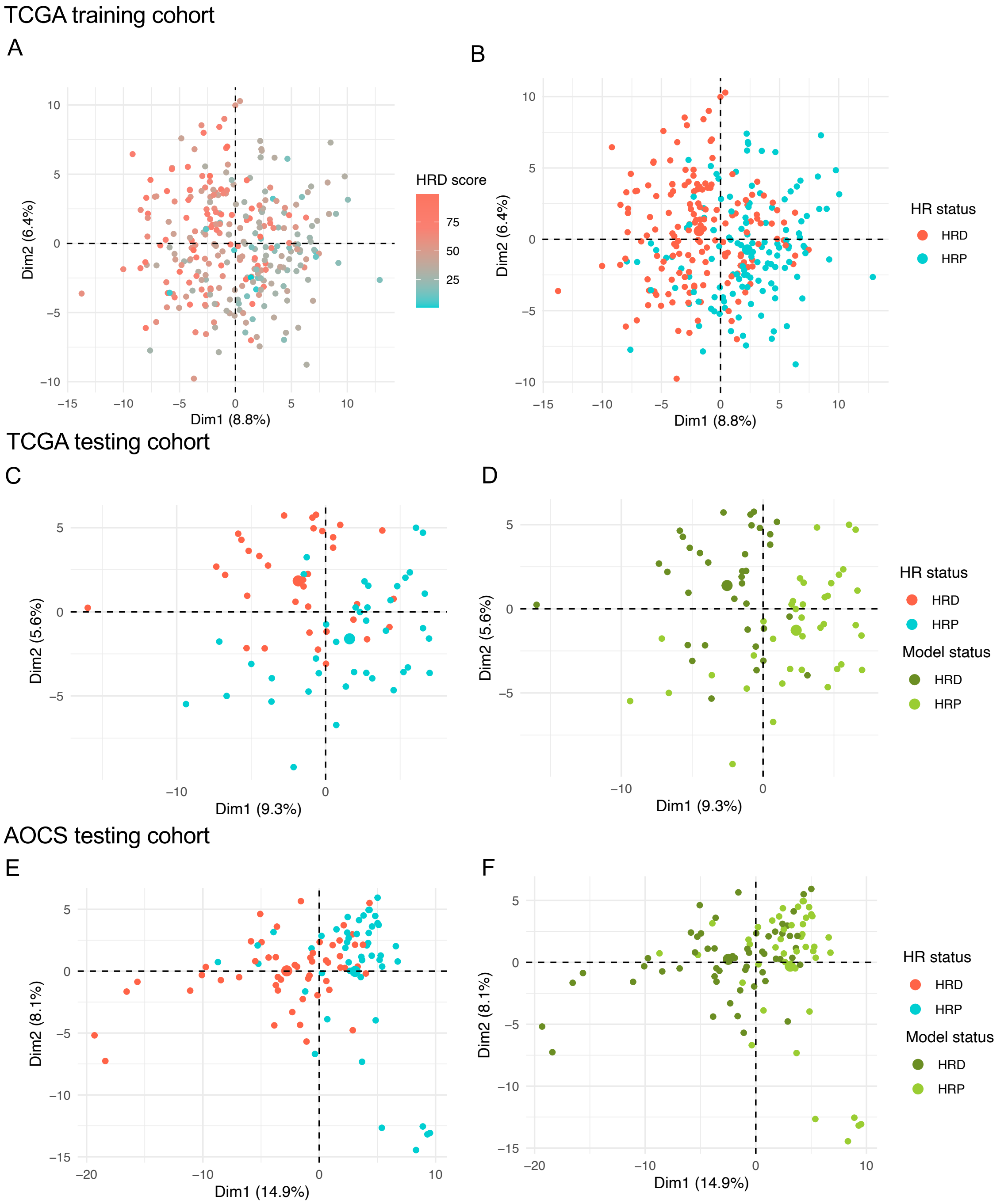


**Supplementary figure 4.** *HGSCs cluster by the normalised and scaled expression of the 209 genes used by IdentifiHR to predict HR status*. PCA plots of the first and second principal components after counts of IdentifiHR genes (n = 209) were normalised by log2 counts-per-million and scaled using a z-score. Each point represents a unique sample of (A, B) the TCGA training cohort (n = 288), (C, D) the TCGA testing cohort (n = 73) and (E, F) the AOCS testing cohort (n = 99). Points coloured by (A) HRD score and (B, C, E) HR status (HRD defined as having a HRD score of ≥ 42) and (D, F) HR status as predicted by the model.

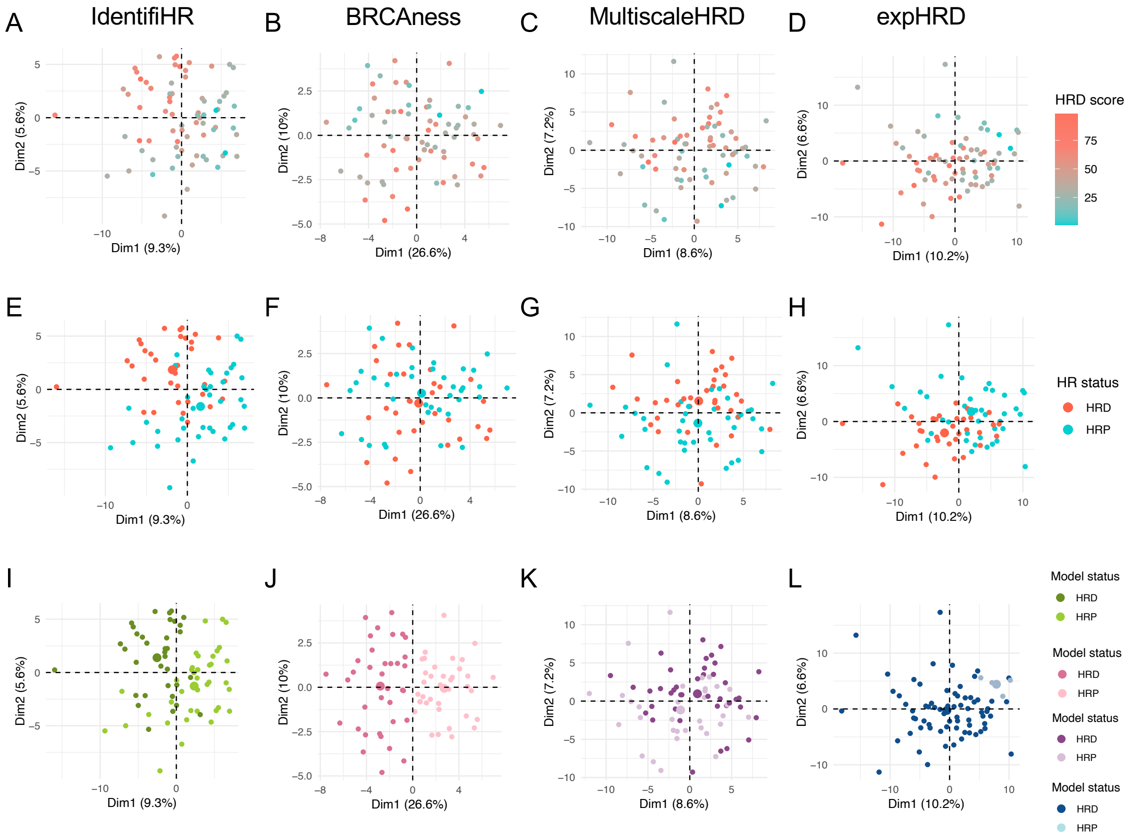


**Supplementary figure 5.** *Qualitatively comparing gene-expression based tools to predict HR status using PCA*. PCA using the normalised and scaled input genes for (A, E, I) IdentifiHR (n = 209 genes), (B, F, J) BRCAness (n = 40 genes), (C, G, K) MultiscaleHRD (n = 228 genes) and (D, H, L) expHRD (n = 356 genes) in the TCGA HGSC testing cohort (n = 73 samples). Samples are clustered by (A, B, C, D) HR status, (E, F, G, H) HRD score and (I, J, K, L) the predicted HR status given by each model.

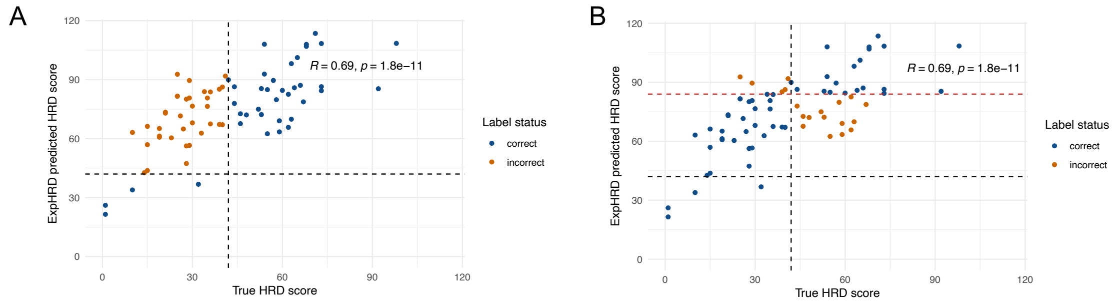


**Supplementary figure 6.** *The relationship between the true HRD score and the HRD score predicted by the expHRD model in the TCGA testing cohort (n = 73 samples)*. The black horizontal and vertical dashed lines at (A) 42, show the clinical HRD score cut-off that separates HRP and HRD and (B) the red horizontal dashed line at 84, to demonstrate the optimal cut-off to maximise model accuracy. Points coloured by whether expHRD predicted the sample's HR status correctly or incorrectly, at each cut-off.
